# Area from image analyses accurately estimates dry-weight biomass of juvenile moss tissue

**DOI:** 10.1101/2020.03.20.000539

**Authors:** Wesley P. Burtscher, Marna A. List, Adam C. Payton, Stuart F. McDaniel, Sarah B. Carey

## Abstract

- Mosses have long served as models for studying many areas of plant biology. Investigators have used two-dimensional measurements of juvenile growth from photographs as a surrogate for dry-weight biomass. The relationship between area and biomass, however, has not been critically evaluated.
- Here we grew axenic tissue cultures of ten *Ceratodon purpureus* isolates to study the relationship between these parameters. We measured area and biomass on replicate cultures with two distinct starting inoculum sizes each week for three weeks and examined the correlation between area and biomass as well as the influence of variation in inoculum size on both parameters.
- We found a strong correlation between area and biomass after two weeks of growth. Furthermore, we found inoculum size affected biomass during the first week of growth but not subsequent weeks and inoculum size had no detectible effect on area.
- These analyses provide experimental confirmation that area is a suitable proxy for biomass and provide clear guidelines for when inoculum size variation may affect downstream growth estimates.

## Introduction

Mosses have long served as models for studying many problems in plant biology (Cove, 2005; Prigge & Bezanilla, 2010). One of the reasons for their utility is the juvenile moss tissue, protonema, is easy to clonally propagate in sterile conditions. Protonema grows in filaments comprised of two cell types (chloronema and caulonema) that divide by serial extension (i.e., along one plane) and frequently branch, forming a dense three-dimensional network. The maturation of protonema triggers the production of buds, which begin to divide along multiple planes, producing mature, leafy gametophores. It is relatively simple to generate axenic protonemal tissue from germinating spores, wounded mature gametophore tissue, or by subsampling or blending growing protonema (Cove *et al*., 2009a). The cell walls of protonema can also be digested to generate abundant protoplasts (Cove *et al*., 2009b). In addition, protonema are responsive to hormone treatments, as well as other chemical or environmental perturbations and are amenable to UV and chemical mutagenesis (Cove, *et al*., 2009c). The fact that identical haploid clonal replicates can be exposed to the same experimental manipulations in a relatively small amount of space, like a growth chamber, makes moss protonema particularly useful.

As a consequence of these attributes, many investigators have used protonemal growth rate as a focal phenotype or a surrogate for fitness (McDaniel *et al*., 2008; Nomura & Hasezawa, 2011; Proust *et al*., 2011; Tani *et al*., 2011; Cho *et al*., 2012; Rawat *et al*., 2017). The standard practice is to photograph protonemal tissue growing on agar-containing Petri dishes under different treatments and measure the area occupied. This approach, however, uses a two-dimensional representation of a three-dimensional phenotype, because protonema grow orthogonal to the surface of the agar (i.e., up) as well as laterally. Thus, two genotypes with very different growth patterns (primarily horizontal growth versus primarily vertical growth) could be scored as different using the typical image analysis procedure. Such variation in growth form is not uncommon among natural isolates or mutagenized plants (McDaniel *et al*., 2008; Perroud & Quatrano, 2008). In other plant systems, dry-weight biomass provides a better estimate of overall growth than does two-dimensional area. Measuring protonema using dry-weight biomass, however, cannot be easily automated, making it a time-consuming procedure, and it can only be measured in effectively dead cells, which negates some of the benefits of working with moss protonema.

In this paper we describe a critical evaluation of the relationship between the two-dimensional area of protonema, estimated from photographs, to the more biologically relevant trait of biomass. We grew ten diverse isolates of the model moss *Ceratodon purpureus* and measured both two-dimensional area and biomass at three time points. We used isolates representing the global diversity of *C. purpureus* because the species is highly polymorphic in growth patterns (Shaw & Beer, 1999; McDaniel, 2005). We found a strong correlation between area and biomass after two weeks of growth. We also varied the starting inoculum size to evaluate how sensitive the two measures of protonemal growth were to such variation. This measure may be important in labs where multiple individuals initiate experimental cultures or starting inoculum size varies because of other experimental variables. We found while we could detect an effect of inoculum size on biomass after one week, we found no effect after two weeks. Moreover, we detected no effect of inoculum size on area after only one week. Collectively these data indicate that area is a suitable proxy for dry-weight biomass and that for *C. purpureus* growth experiments the effect of inoculum size variation disappears after two weeks.

## Materials and Methods

### Tissue generation

The *C. purpureus* isolates used in this study were previously grown from single-spore isolates and maintained as lab cultures. We used five male/female sibling pairs (i.e., a male and female from the same sporophyte) that were originally isolated from Ecuador, Chile, New York, Connecticut, and Alaska. To generate tissue for testing area against biomass and the effect of starting inoculum size, we grew axenic tissue cultures following Cove *et al*., 2009a. We divided the plates into two inoculum treatments: small and large. For the small inoculum size, the moss tissue was rolled into balls that were small but visually uniform in initial size. Each time point had 15 Petri dishes containing four isolates (although a few only had two isolates). Each isolate was replicated six times per time point and to which plate each replicate was placed was randomly selected. For the large inoculum size, the moss tissue was rolled into larger balls of visually uniform initial size (approximately twice the small inoculum size). Each time point had eight Petri dishes of four randomly selected isolates (again a few only had two isolates). Three clonal replicates of each isolate were used per time point for the large inoculum size analyses and to which plate each isolate was grown was randomly selected. The same tissue for the inoculum-size analyses was used for comparisons of area and biomass. The tissue was grown on cellulose cellophane disks overlaying 0.7% agar (A9799, Plant Cell Culture Agar; Sigma, St Louis, MO, USA) in 100 mm Petri dishes containing BCD medium and 5 mM (di)ammonium tartrate (Cove *et al*., 2009a). Cultures were grown at a constant temperature of 25°C in 18 hour full light days in Percival Scientific growth chambers (Percival Scientific, Inc., Perry, Iowa) with a light intensity 60–80 μmol m^−2^ s^−1^. We haphazardly rotated plate positions inside the growth chamber every 24 hours.

### Data collection

Measurements of area and biomass and of small and large inoculum sizes were taken at three time points: at one, two, and three weeks of growth. To collect area measurements, each plate was photographed using a Canon EOS 50D camera (Canon, Inc., Ota City, Tokyo, Japan) and assessed using ImageJ (Schneider *et al*., 2012). ImageJ measures area by converting pixels to mm^2^ using an object of known length present in the photographs. To collect biomass results, at each time point we harvested the designated replicates and placed each separately into a 1.5 mL tube of known weight (which we measured beforehand). The tissue was then dried in a precision gravity convection incubator for four days at 37°C. We weighed the dried tissue to the 0.1 mg accuracy on a Mettler Toledo scale (Mettler Toledo, Columbus, Ohio).

The final data set included 264 replicates—between 25 and 27 for each of ten isolates. We removed two clonal replicates that died; several others showed signs of bacterial contamination, but we have included these in our analyses.

### Statistical analyses

All statistical analyses and plots were done in R (version 3.5.3; R Core Team, 2019). To test whether area, a two-dimensional representation of moss growth, is an acceptable proxy for biomass we first averaged biomass and area for all clonal replicates within each inoculum size for each single-spore isolate and ran separate correlations for each week using cor().To test whether starting inoculum size of protonemal tissue affects the results of area and biomass measurements, we ran ANOVAs on each week separately using aov().

## Results

To test the strength of the relationship between change in area and change in biomass, we ran a correlation of area by biomass for each week’s data. We found a weak correlation between area and biomass for week one (0.389; Fig. **1a**) but strong correlations between the two variables in weeks two and three (0.858 and 0.856, respectively; Fig. **1b,c**).

**Figure 1.**
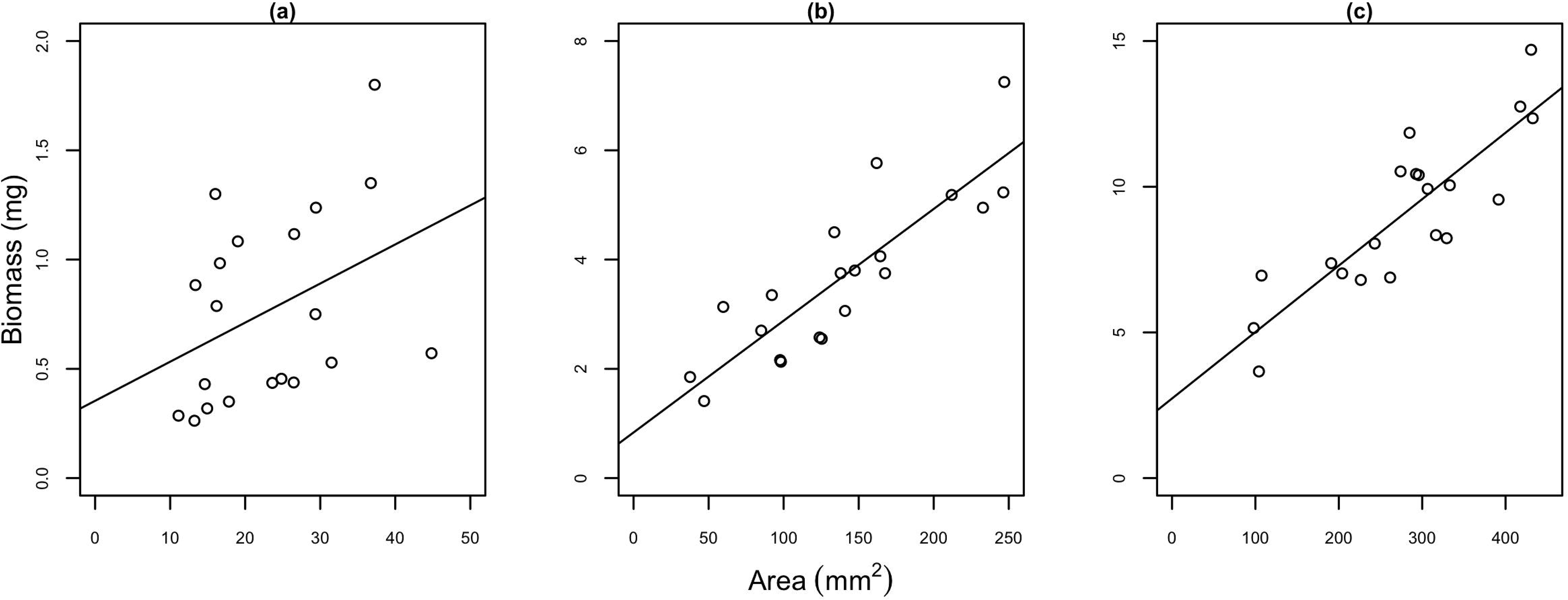
Scatterplots of mean biomass (mg) and mean area (mm^2^) during three weeks of growth in large and small inoculum sizes of ten isolates of *Ceratodon purpureus*. Area and biomass were compared at **(a)** one, **(b)** two, and **(c)** three weeks of growth. Week one shows a weak positive correlation (0.393), while weeks two (0.858) and three (0.837) show a strong positive correlation. Lines are linear least squares regression fits to data points.

To test the effect of inoculum size on area and biomass, we used an ANOVA to compare either area or biomass with inoculum size for each of the three weeks. Inoculum size had a detectable effect on biomass in week one (F_1,18_ = 28.51; p = 4.48×10^−5^; Fig. **2a**) but no effect in weeks two (F_1,18_ = 2.139; p = 0.161; Fig. **2b**) or three (F_1,18_ = 1.127; p = 0.302; Fig. **2c**). In contrast, inoculum size had no effect on area at any week measured (week 1 F_1,18_ = 0.038; p = 0.847; week 2 F_1,18_ = 0.208; p = 0.654; week 3 F_1,18_ = 0.088; p = 0.77; Fig. **3a,b,c**).

**Figure 2.**
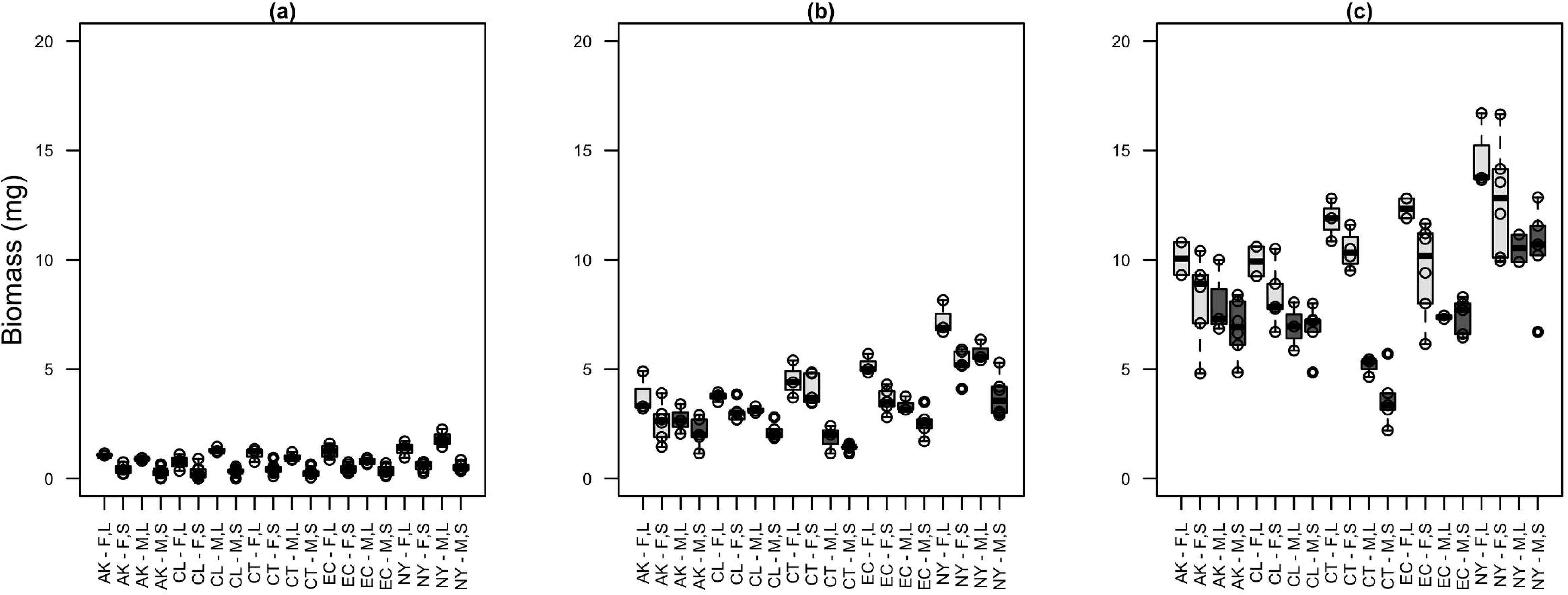
Boxplots of biomass (mg) of *C. purpureus* isolates after one (a), two (b), and three (c) weeks of growth. Different populations (AK = Alaska, CL = Chile, CT = Connecticut, EC = Ecuador, NY = New York) of female (F) and male (M) isolates at large (L) and small (S) inoculum sizes have a significant difference in biomass at one week **(a)** but show no significant difference in weeks two and three **(b)** and **(c)**.

**Figure 3.**
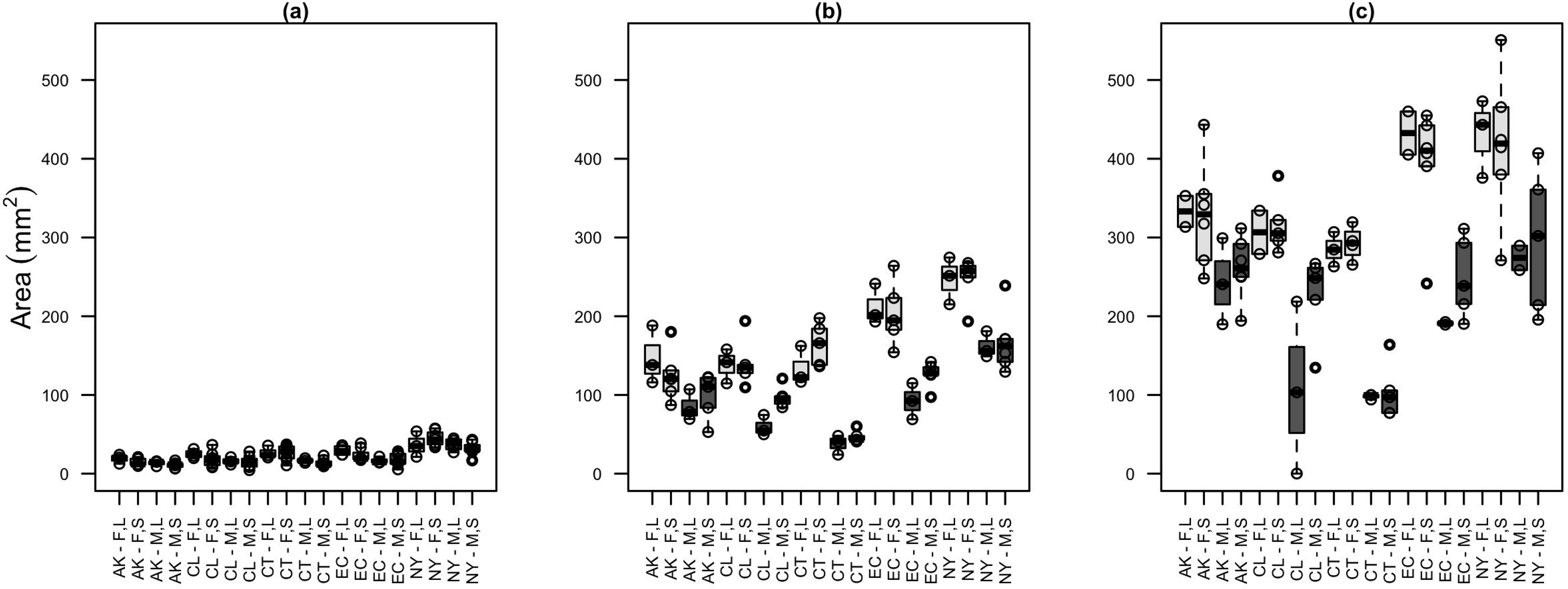
Boxplots of area (mm^2^) of isolates after one **(a)**, two **(b)**, and three **(c)** weeks of growth. Different populations (AK = Alaska, CL = Chile, CT = Connecticut, EC = Ecuador, NY = New York) of female (F) and male (M) isolates at large (L) and small (S) inoculum sizes have no significant difference in area at any of the three weeks tested.

## Discussion

It is common practice to use image analysis of two-dimensional area as an estimate of growth or fitness in moss protonema. This procedure is non-invasive, fast, and easy to automate. Nevertheless, the accuracy with which this process estimates biomass, the underlying biologically relevant value for assessing overall growth, has not been critically evaluated. Here we show two-dimensional area can provide a strong approximation for protonemal biomass, with some caveats. We found a weak but positive correlation between area and biomass after one week of growth. However, the correlation between these parameters was very strong and stable after two weeks of growth. In addition, the initial size of each replicate (inoculum size) had a negligible effect on that growth. We did find that inoculum size had an effect on biomass at one week of growth, possibly because either we were very close to the range of detection of our scale or the smaller inocula took more time to initiate growth. However, we found no significant difference after two weeks. Additionally, we found inoculum size had no detectable effect on protonemal area, meaning area may be a less-biased measure of growth than biomass under some circumstances. Therefore, while it is still recommended to make moss inocula that are reasonably close in size, concerns about starting tissue affecting downstream growth results are unnecessary (at least within inoculum sizes where the largest is about twice the size of the smallest).

Our tissue culture work with other mosses suggests the relationship between two-dimensional area and biomass is likely to hold true across this group. The growth architecture of *Physcomitrella patens*, the most widely used moss model, is quite similar to *C. purpureus* at the protonemal stage, in spite of the fact that these species last shared a common ancestor more than 200 million years ago (Laenen *et al*., 2014). Our experience growing other mosses more closely related to *C. purpureus* suggests this growth pattern is widely conserved. The trends and associations are also consistent between males and females, despite the obvious sexual dimorphism in protonemal growth rate in this species (McDaniel *et al*., 2008; Fig. **2**,**3**). However, in some mosses, such as *Aulacomnium palustre*, the cultures produce limited protonemal growth and develop gametophores much faster than in either *C. purpureus* or *P. patens*. Such species may require additional care when interpreting area as a proxy for growth. Similarly, some environmental conditions may promote growth forms that are more likely to introduce error in the method that we describe. Elevated humidity, for example, induces vertical growth in some genotypes, and media with decreased calcium concentrations can induce a very compact growth. However, under most circumstances in standard growth conditions, if environmental conditions are consistent across samples, two-dimensional area is likely to provide an accurate estimate of biomass and therefore growth. Thus, taken together these results will be of use for design and implementation of moss protonemal tissue growth analyses, as well as tissue with a similar growth phenotype.

## Acknowledgments

This work was funded by a University of Florida University Scholars Program fellowship to M.A.L., a University of Florida Department of Biology Mildred Mason Griffith Botany Grant to S.B.C., and an NSF grant to S.F.M. (DEB 1541005).

## Author Contributions

This study was conceived by A.C.P., M.A.L., S.B.C., and S.F.M. The growth experiment was set up by M.A.L. and photograph collection was by A.C.P., M.A.L., and S.B.C. ImageJ analyses were done by M.A.L. Statistical analyses were conducted by S.B.C and W.P.B. Interpretation of results and writing of the manuscript were done by M.A.L., S.B.C., S.F.M and W.P.B.

## Supporting Information

Table S1. Data for area and biomass statistical analyses and scatterplot, where replicates within an isolate, inoculum size, and week were averaged.

Table S2. Data for boxplots of area and biomass, where replicates within an isolate, inoculum size, and week were not averaged.

## References

Cho, SH, Ceyda C, Axtell MJ. 2012. miR156 and miR390 regulate tasiRNA accumulation and developmental timing in *Physcomitrella patens*. The Plant Cell 24.12 (2012): 4837–4849.

Cove D. 2005. The Moss Physcomitrella patens. Annu. Rev. Genet. 39: 339–358. 10.1146/annurev.genet.39.073003.110214.

Cove DJ, Perroud P, Charron AJ, McDaniel SF, Khandelwal A, and Quatrano RS. 2009a. Isolation and regeneration of protoplasts of the moss *Physcomitrella patens*. Cold Spring Harbor Protocols 2009 2: pdb.prot5140.

Cove D, Perroud P-F, Charron A, McDaniel S, Khandelwal A, Quatrano R. 2009b. The moss *Physcomitrella patens*: a novel model system for plant development and genomic studies. Emerging Model Organisms. New York, NY, USA: Cold Spring Harbor Laboratory Press.

Cove DJ, Perroud P, Charron AJ, McDaniel SF, Khandelwal A, and Quatrano RS. 2009c. Transformation of the moss *Physcomitrella patens* using direct DNA uptake by protoplasts. Cold Spring Harbor Protocols 2009 2: pdb.prot5143.

Laenen B, Shaw B, Schneider H, Goffinet B, Paradis E, Désamoré A, Heinrichs J, Villarreal JC, Gradstein SR, McDaniel SF et al. 2014. Extant diversity of bryophytes emerged from successive post-Mesozoic diversification bursts. Nature Communications 5: 6134.

McDaniel SF. 2005. genetic correlations do not constrain the evolution of sexual dimorphism in the moss *Ceratodon purpureus*. Evolution 59: 2353–2361. 10.1111/j.0014-3820.2005.tb00945.x.

McDaniel SF, Willis JH, Shaw AJ. 2008. The genetic basis of abnormal development in interpopulation hybrids of the moss *Ceratodon purpureus*. Genetics 179: 1425–1435.

Nomura, T., Hasezawa, S. Regulation of gemma formation in the copper moss *Scopelophila cataractae* by environmental copper concentrations. J Plant Res 124, 631–638 (2011). https://doi.org/10.1007/s10265-010-0389-3

Prigge MJ, Bezanilla M. 2010. Evolutionary crossroads in developmental biology: *Physcomitrella patens*. Development 137: 3535–3543.

Proust H, Hoffmann B, Xie X, Yoneyama Kaori, Schaefer DG, Yoneyama K, Nogué F, Rameau C. 2011. Strigolactones regulate protonema branching and act as a quorum sensing-like signal in the moss *Physcomitrella patens*. Development 138: 1531–1539.

R Core Team. 2019. R: a language and environment for statistical computing. Vienna, Austria: R Foundation for Statistical Computing. http://www.R-project.org/

Rawat A, Brejšková L, Hála M, Cvrčková F, and Žárský V. 2017. The Physcomitrella patens exocyst subunit EXO70.3d has distinct roles in growth and development, and is essential for completion of the moss life cycle. New Phytol. 216: 438–454. 10.1111/nph.14548.

Schneider CA, Rasband WS, Eliceiri KW. 2012. NIH image to ImageJ: 25 years of image analysis. Nature Methods 9: 671–675.

Shaw AJ, Beer SC. 1999. Life history variation in gametophyte populations of the moss *Ceratodon purpureus* (Ditrichaceae). American Journal of Botany 86: 512–521.

Tani A, Akita M, Murase H, and Kimbara K. 2011. Culturable bacteria in hydroponic cultures of moss Racomitrium japonicum and their potential as biofertilizers for moss production. Journal of Bioscience and Bioengineering 112 1:32–39. http://www.sciencedirect.com/science/article/pii/S1389172311001162.

